# A stress-function tradeoff organizes epithelial heterogeneity across spatial scales in the human thyroid

**DOI:** 10.64898/2026.03.12.711294

**Authors:** Yael Korem Kohanim, Tal Barkai, Roy Novoselsky, Sapir Shir, Keren Bahar Halpern, Shlomit Reich-Zeliger, Jacob Elkahal, Idit Tessler, Shaked Shivatzki, Ignat Schwartz, Eric Remer, Galit Avior, Rouven Hoefllin, Merav Kedmi, Hadas Keren-Shaul, Inna Goliand, Yoseph Addadi, Ofra Golani, Eran Alon, Shalev Itzkovitz, Ruslan Medzhitov

## Abstract

Many organs are organized into repeating anatomical units, yet how cellular heterogeneity is structured within and between these units remains poorly understood. Here we use spatial transcriptomics to dissect multiscale heterogeneity in the human thyroid gland, a tissue composed of hormone-producing follicles. Across human thyroid samples spanning non-inflamed to inflamed states, we develop a follicle-aware analytical framework that separates intra-follicular from inter-follicular variability. We find that heterogeneity among thyrocytes is not dominated by differences in hormone synthesis but instead by two opposing transcriptional programs: an active hormone-producing state and a damage-response thyrocyte (DRT) state enriched for stress, immune, and damage-response pathways. DRTs are spatially clustered, associated with DNA damage markers, and are enriched near immune niches. Notably, the balance between active and damage-response programs constitutes a major axis of variability across cells, follicles, and patients. Our findings highlight a damage-response epithelial thyrocyte state that may be fundamental to follicular function in the human thyroid and provide a general framework for studying heterogeneity in tissues composed of repeating anatomical units.

## Introduction

Tissues in the human body often achieve complex physiological functions through the repetition of autonomous anatomical units^1^. Examples include thyroid follicles, intestinal villi, and liver lobules. While this organizational principle is central to tissue function, how cellular heterogeneity is structured within and between such repeating units remains poorly understood. In particular, it is unclear whether variability primarily reflects differences in core functional outputs, stochastic noise, or adaptive responses to local microenvironmental stress^2^.

Recent advances in spatial transcriptomics have enabled systematic mapping of gene expression in intact tissues while preserving spatial context ^1,3–8^. These approaches have begun to reveal previously unappreciated cellular diversity and spatial organization in both healthy and diseased tissues. However, most analyses have focused on global variations along the major axes of stereotypical organ units, often ignoring differences and potential division of labor between individual units. Dissecting variability between units could expose important principles of tissue-level organization.

The human thyroid gland provides a powerful system to address the questions of sources of heterogeneity within and between repeating anatomical units. The thyroid gland is composed of follicles – discrete, spherical units formed by monolayers of thyrocytes surrounding a colloid-filled lumen – which together are responsible for the synthesis and secretion of thyroid hormones. Although thyrocytes are traditionally viewed as functionally homogeneous within follicles, thyroid hormone synthesis requires the coordination of multiple metabolically demanding steps, which may trade off with one another and give rise to heterogeneous gene expression patterns^9,10^. Moreover, sustained hormone production imposes substantial metabolic and oxidative stress, exposing follicular cells to elevated levels of reactive oxygen species. In addition, thyroid tissue is frequently exposed to autoimmunity, with autoimmune thyroid diseases being the most common autoimmune disorders^11^. Approximately 20% of the population carries autoantibodies against thyroid antigens^12^, and even samples from individuals without overt thyroid disease often exhibit subclinical immune infiltration^13^. Together, these features raise the possibility that thyrocytes may adopt distinct functional states to balance hormone production with tissue maintenance and stress adaptation.

Previous single-cell transcriptomic studies have identified heterogeneity among thyrocytes, including stress- and immune-associated programs, but have largely lacked spatial resolution or explicit follicular context^14–18^. Consequently, it remains unclear how such states are distributed within follicles, whether they vary systematically between follicles, and how they relate to local immune activity. Addressing these questions requires an analytical framework that explicitly integrates spatial information with the hierarchical organization of thyroid tissue.

Here, we use spatial transcriptomics at multiple resolutions to dissect cellular heterogeneity in human thyroid tissue across health and inflammation. By combining cell-and follicle-level segmentation with a multinomial randomization framework, we develop a follicle-aware approach that separates intra-follicular from inter-follicular variability. Applying this framework to ten human thyroid samples, we uncover a damage-response thyrocyte state that emerges as a dominant axis of heterogeneity across spatial scales. We show that this state is associated with stress and DNA damage responses, is spatially clustered within and between follicles, enriched near immune niches, and distinct from previously described MHC class II-expressing thyrocytes. Furthermore, we show that the balance between active and damage-response thyrocyte programs defines a major axis of inter-patient variability that shifts with age. Together, our findings reveal how stress-associated epithelial states organize follicular function in the human thyroid gland and provide a general framework for studying multiscale heterogeneity in tissues composed of repeating anatomical units.

## Results

### Spatial transcriptomics enables multiscale analysis of thyrocyte heterogeneity in human thyroid tissue

To investigate how cellular heterogeneity is organized within the thyroid gland, we analyzed human thyroid samples using spatial transcriptomics across a continuum of histological states ranging from non-inflamed to inflamed tissue (Fig. 1a, Table S1). We profiled ten human thyroid samples using both 10x Genomics Visium (n=8) and the higher-resolution VisiumHD platforms (n=2), enabling complementary interrogation of inter-follicular and intra-follicular variation. To explicitly capture tissue organization, we used machine learning-based segmentation of the histological images to delineate individual cells and follicles (Fig. 1b-d, Supplementary Fig. 1, Methods). VisiumHD spatial bins were aggregated to enable single-cell level analysis, and individual cells were assigned to their corresponding follicles, resulting in a hierarchical data structure linking cellular and follicular gene expression (Fig. 1c,d). This framework enabled separation of heterogeneity occurring among cells within the same follicle (intra-follicular variability) from heterogeneity between distinct follicles (inter-follicular variability), providing a principled approach to studying functional variation in a tissue composed of repeating anatomical units (Fig. 1e).

**Figure 1.**
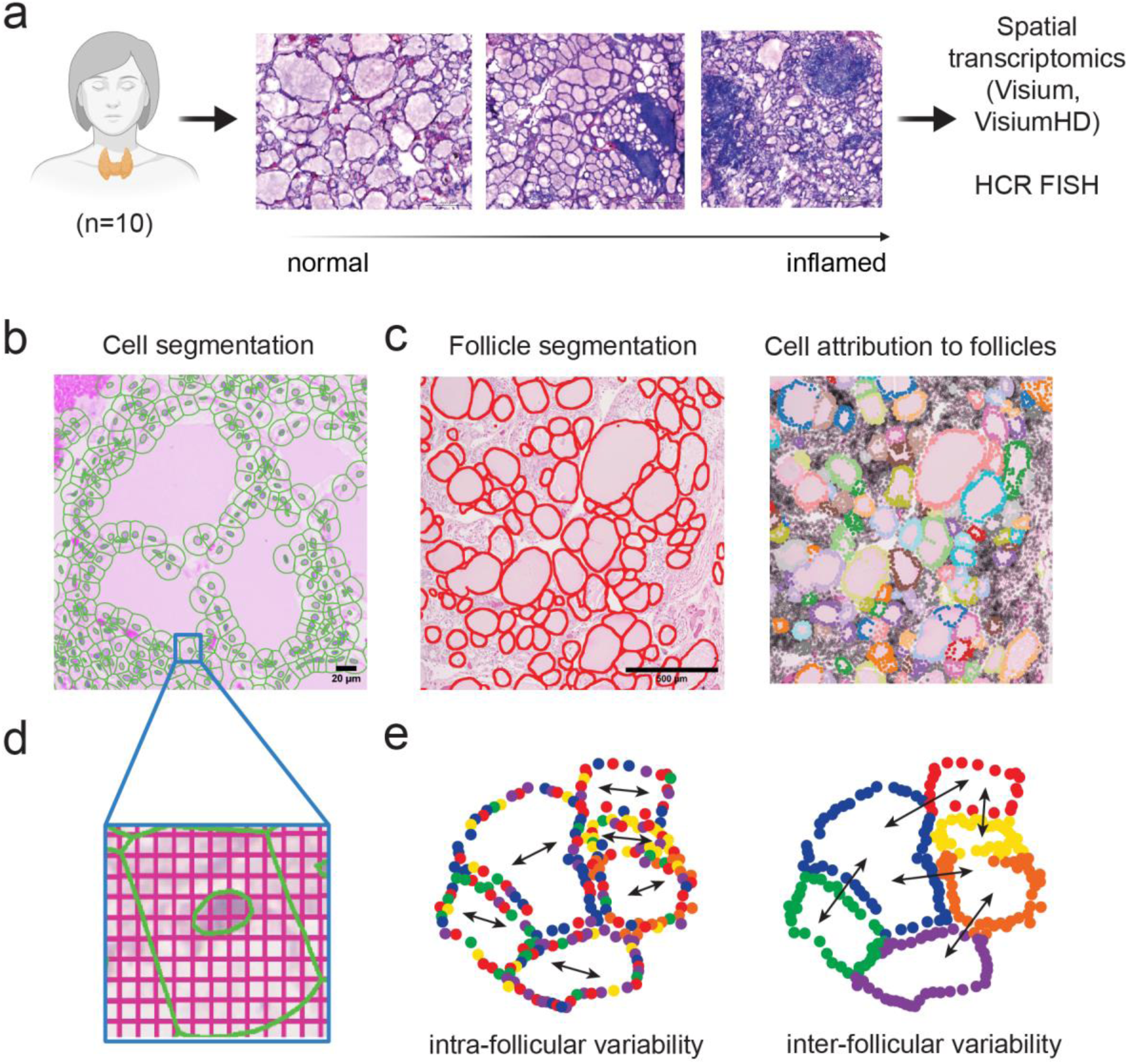
Spatial transcriptomics framework enables multiscale analysis of thyrocyte heterogeneity. **a**, Study overview: human thyroid samples (n=10) spanning non-inflamed to inflamed states profiled by spatial transcriptomics (Visium, VisiumHD) with hybridization chain reaction fluorescence *in situ* hybridization (HCR FISH) validation^19^. **b**, Cell segmentation of histological images. **c**, Follicle segmentation (left) and assignment of cells to follicles (right, cells are randomly colored according to follicle assignment, unassigned cells are in gray). **d**, VisiumHD 2×2μm bins were summed according to cell segmentation to create an aggregated single-cell structure. **e**, Conceptual framework illustrating separation of intra-follicular and inter-follicular variability enabled by follicle-aware spatial analysis.

### Intra-follicular variability is dominated by two opposing functional programs

We first asked whether thyrocytes within the same follicle exhibit transcriptional variability. For each thyrocyte-specific gene, we quantified intra-follicular variability by computing the standard deviation (SD) of expression across thyrocytes within individual follicles (Methods). To determine whether observed variability exceeded technical and sampling noise, we compared these values to a null distribution generated by multinomial randomization, preserving both the total number of unique molecular identifiers (UMIs) per cell and the total number of UMIs per gene within each follicle (Fig. 2a, Methods).

**Figure 2.**
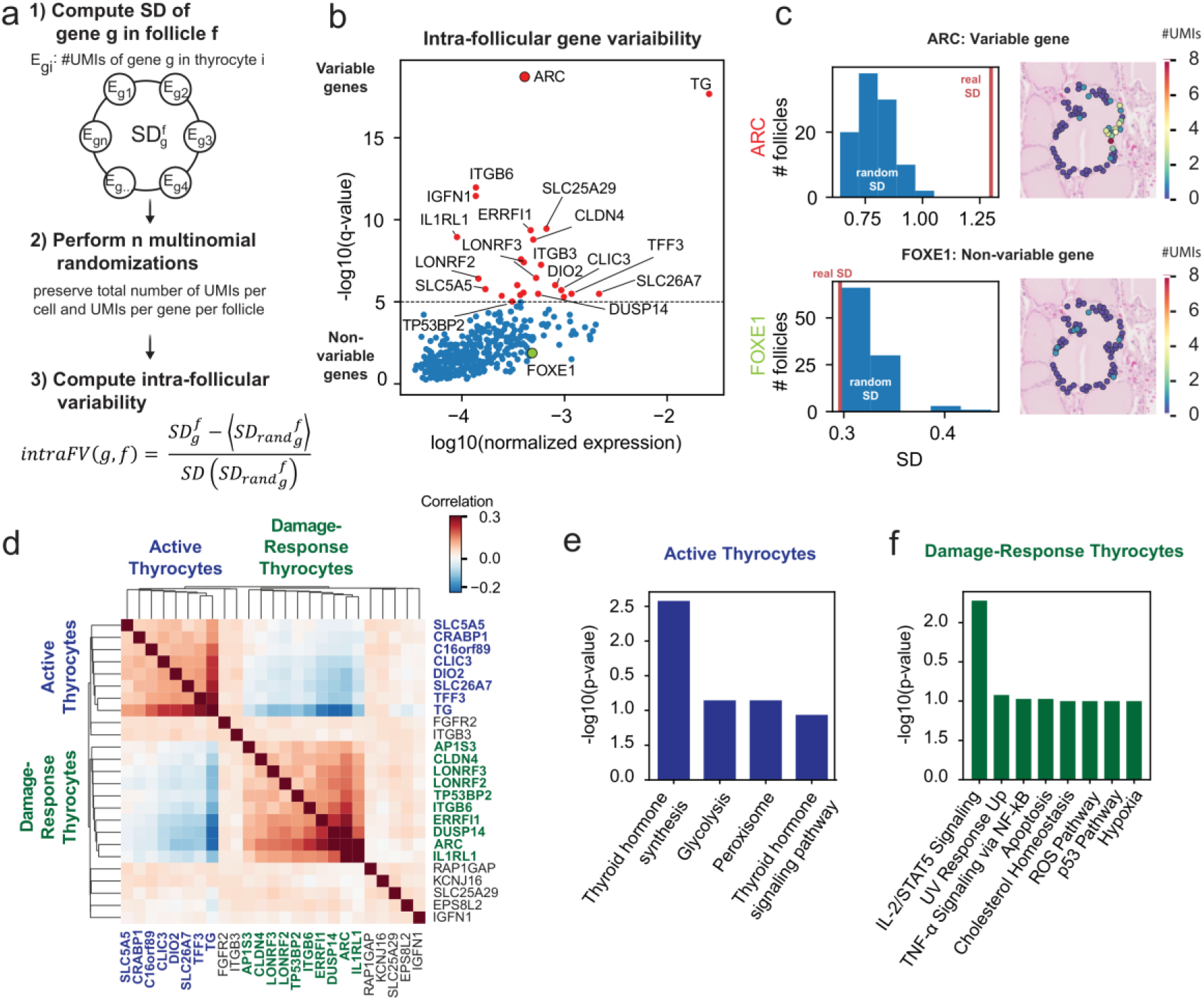
Intra-follicular variability reveals two opposing functional gene expression modules. **a**, Multinomial randomization framework to quantify intra-follicular variability while preserving total UMIs per cell and per gene within each follicle. The standard deviation (SD) of the number of UMIs of gene *g* is computed for each follicle and compared to randomized follicles (Methods). **b**, Intra-follicular variability analysis highlights a subset of thyrocyte genes that are variable within follicles. Q-values are the adjusted p-values of the gene standard deviation based on the randomization tests. **c**, Example distributions comparing observed versus randomized standard deviations for a variable gene (*ARC*, colored red with black outline, in panel b) and a non-variable gene (*FOXE1*, colored green in panel b) in a follicle with similar expression of these genes, with corresponding spatial expression maps. Expression levels are UMI counts **d**, Spearman correlation matrix of genes with significant intra-follicular variability across single cells in VisiumHD (q-value<10^−5^), revealing two anti-correlated gene modules: the Active thyrocyte module (blue) and the Damage-response thyrocyte (DRT) module (green). Significantly variable genes were defined by high intra-follicular variability. Correlation values were trimmed at *r*=0.3, values on the diagonal are *r*=1. **e,f**, Pathway enrichment analysis of the active thyrocyte module **(e)** and DRT module **(f)**.

This analysis revealed that only a restricted subset of genes displayed significantly elevated intra-follicular variability relative to randomized expectations (Fig. 2b, Table S2). For example, the gene *ARC* exhibited gene expression variability that significantly exceeded that expected at random, whereas the gene *FOXE1* showed expression variability within the expected range of the random distribution (Fig. 2c).

Next, we asked whether the intra-follicular variable genes are arranged in coherent expression modules. Indeed, hierarchical clustering of the correlation matrix of genes that exhibited significant intra-follicular variability uncovered two strongly anti-correlated expression programs (Fig. 2d). One program was enriched for genes involved in thyroid hormone synthesis and iodide metabolism, such as thyroid hormone precursor thyroglobulin (*TG*), iodide transporters *SLC5A5* (Sodium/iodide cotransporter, NIS) and *SLC26A7*, and iodothyronine deiodinase 2 (*DIO2*), consistent with an active hormone-producing state (Fig. 2d,e). The second program was enriched for stress response, immune signaling, and damage-associated pathways, defining a damage-response transcriptional program (Fig. 2d,f). Pathway enrichment analysis highlighted multiple stress pathways in DRT thyrocytes, including Reactive Oxygen Species, Hypoxia, P53 and TNF-alpha signaling (Fig. 2f)^20^. Two prominent genes in this module were the canonical IL33 receptor *IL1RL1*, and *ARC*, a gene that has been studied mainly in the context of neurons^21,22^. We validated the co-expression of these genes by hybridization chain reaction fluorescence *in situ* hybridization (HCR FISH) (Supplementary Fig. 2). We observed an anti-correlation between the active and damage-response modules in both Visium and VisiumHD platforms (Supplementary Fig. 3). These results indicate that intra-follicular heterogeneity among thyrocytes is dominated by a balance between active metabolic function and a distinct damage-response state.

### Damage-Response Thyrocytes shape heterogeneity across follicles

Early physiological and histological studies have suggested that functional differences between thyrocytes can be amplified at the level of individual follicles and across the gland^23,24^. We therefore next asked which programs shape variability between follicles (Fig. 3a). To quantify inter-follicular variability, gene expression was aggregated at the follicular level and analyzed using a multinomial randomization framework that preserves total UMIs per follicle and per gene, analogous to that used for intra-follicular variability (Fig. 3a, Methods). We identified 181 genes exhibiting inter-follicular variability that is higher than that expected at random (q-value<0.05) (Table S3). Hierarchical clustering of the most variable genes revealed two modules of correlated genes (Fig. 3b,c). Notably, these significantly overlapped the active and DRT gene modules that showed elevated intra-follicular variability (hypergeometric p-value=7×10^−4^ for active, p-value=2×10^−11^ for DRT).

**Figure 3.**
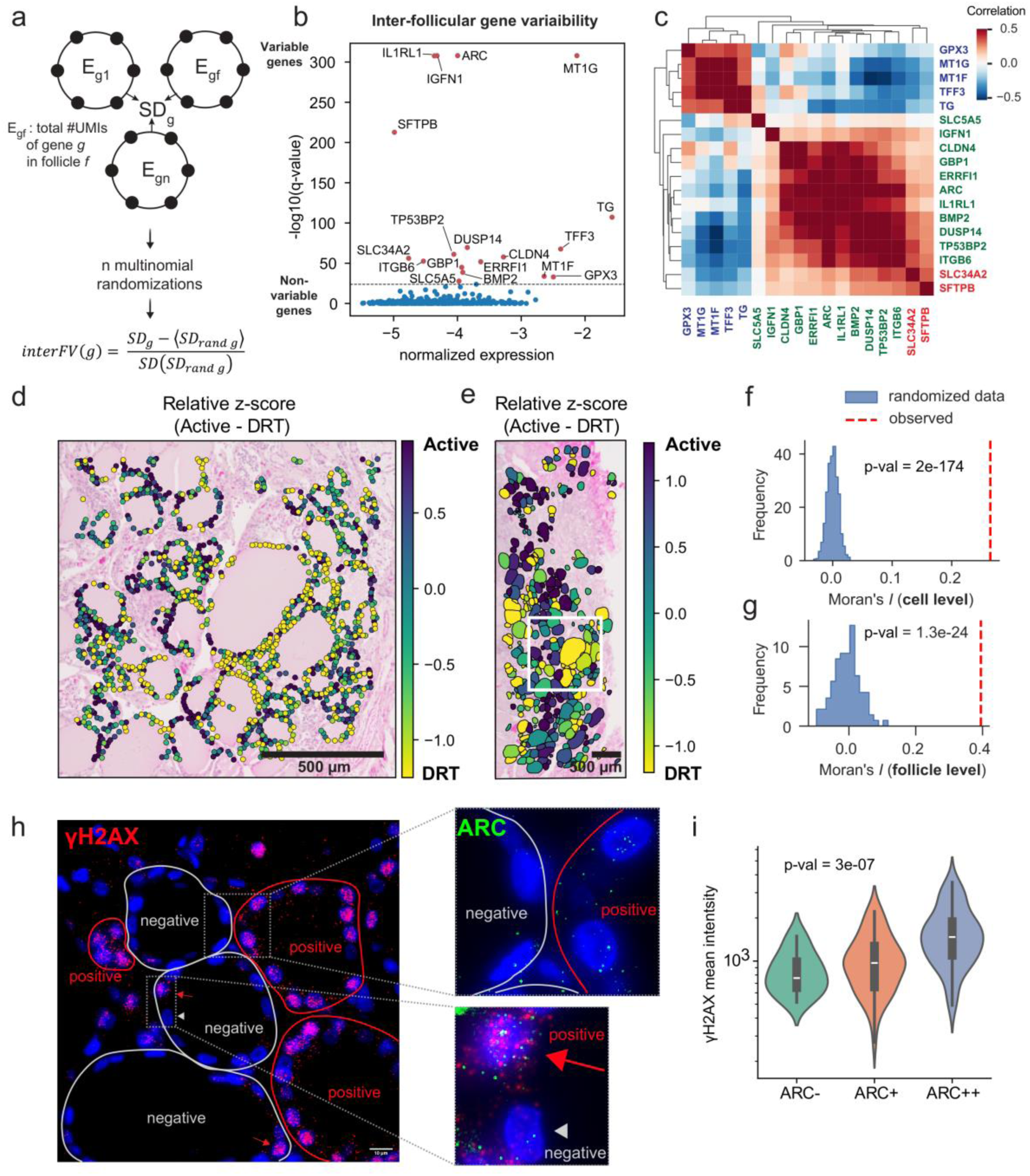
Damage-response thyrocytes shape heterogeneity across spatial scales and associate with DNA damage. **a**, Schematic of inter-follicular variability analysis using multinomial randomizations, preserving total UMIs per follicle and per gene. **b**, Inter-follicular variability significance versus expression for thyrocyte genes. Q-values are the adjusted p-values of the inter-follicle gene standard deviation based on the randomization tests. **c**, Spearman correlation matrix of genes with significant inter-follicular variability across single cells in VisiumHD (q-value<10^−30^), showing module structure at the follicle level. Blue (active) and green (DRT) modules are analogous to intra-follicular modules, additional module in red. Correlation values were trimmed at *r*=0.5, values on the diagonal are *r*=1. **d**, Spatial map of relative z-score difference (Active − DRT) at cellular resolution, illustrating mixed states within follicles. The relative z-score is the difference between the standardized summed expressions of the active module genes and the DRT module genes. **e**, Spatial map of relative z-score highlighting follicular-scale heterogeneity and patchy organization. White box represents the magnified region in panel d. **f**, Moran’s *I* distribution (Methods) at the *cell level* comparing observed spatial autocorrelation to randomized data. **g**, Moran’s *I* distribution at the *follicle level* comparing observed spatial autocorrelation to randomized data. **h**, Combined HCR FISH/HCR IF showing γH2AX (DNA damage marker) and *ARC* signal, illustrating ARC-associated DNA damage response in thyrocytes. Left - γH2AX protein (red dots) displays a patchy spatial pattern. γH2AX positive follicles encircled in red, negative in gray. Scalebar is 10µm. Top right magnified view shows colocalization of ARC mRNA (green dots) with γH2AX in a γH2AX-positive follicle compared with a γH2AX-negative follicle. Bottom right magnified view shows ARC and γH2AX colocalization in a positive versus a negative cell. Red arrows indicate γH2AX-positive cells, gray arrowheads indicate γH2AX-negative cells. **i**, Quantification of γH2AX mean intensity in cells stratified by *ARC* expression (no expression: ARC−; low expression ARC+; high expression ARC++, Methods). Results based on 7 samples from 3 patients (Supplementary Fig. 4b). Box plots white line indicates the median, boxes span the interquartile range (IQR, 25,75 percentiles), whiskers extend up to 1.5 IQR.

We observed a patchy organization of DRTs both within and between follicles. (Fig. 3d,e, Methods). Within a follicle, DRTs tended to cluster together, having DRT neighbors at higher probability than that expected by chance (Fig. 3d,f, Moran’s *I*=0.26, p-value =2×10^−174^). Similarly, follicles with high expression of DRTs tended to have DRT-high neighboring follicles at high probability (Fig. 3e,g Moran’s *I*=0.4, p-value =10^−24^, Methods).

### Damage-response thyrocytes engage in DNA damage response

What could be the functional significance of DRT-high thyrocytes? Thyroid hormone production is associated with the generation of massive amounts of reactive oxygen species (ROS), potentially yielding stressful cellular states^25^. DRT module genes showed elevated levels of ROS and other stress-related pathways (Fig. 2f). To examine whether DRT-high thyrocytes display elevated DNA damage levels, we used HCR FISH of the DRT module marker *ARC*, combined with HCR immunofluorescence (HCR IF) of phosphorylated histone H2AX (γH2AX), a protein that accumulates at DNA double-strand breaks and is engaged in their repair^26^ (Fig. 3h,i, Supplementary Fig. 4a). We found that some follicles showed high levels of γH2AX in most thyrocytes, whereas in other follicles, γH2AX levels were elevated in a minority of thyrocytes (Fig. 3h). Notably, the levels of γH2AX were significantly higher in ARC-high thyrocytes both within and between follicles (Fig. 3i, Supplementary Fig. 4b, p-value=3×10^−7^). This correlated expression of *ARC* in thyrocytes that display high levels of DNA damage response suggests that DRTs may be engaged in DNA repair and stress management.

### Damage-response thyrocytes localize near immune niches and are distinct from MHC class II-expressing thyrocytes

Given the enrichment of immune- and stress-related pathways in damage-response thyrocytes, we next investigated their spatial relationship to immune zones defined by lymphocyte aggregates (Fig. 4, Supplementary Fig. 5). To this end, we examined how DRT-module and active-module summed gene expression varied with distance to immune zones (Fig. 4a,b, Supplementary Fig. 5). Notably, in samples exhibiting immune cell infiltration, DRTs were enriched near immune zones (Fig. 4b,e – Spearman correlation of DRT module gene expression sum to immune zones *r*=-0.14, p-value=2×10^−10^, Supplementary Fig. 5c,f). Conversely, the active thyrocyte state was depleted in proximity to immune regions (Fig. 4a,d, Spearman correlation with distance to immune zones *r*=0.25, p-value=3×10^−29^, Supplementary Fig. 5b,e).

**Figure 4.**
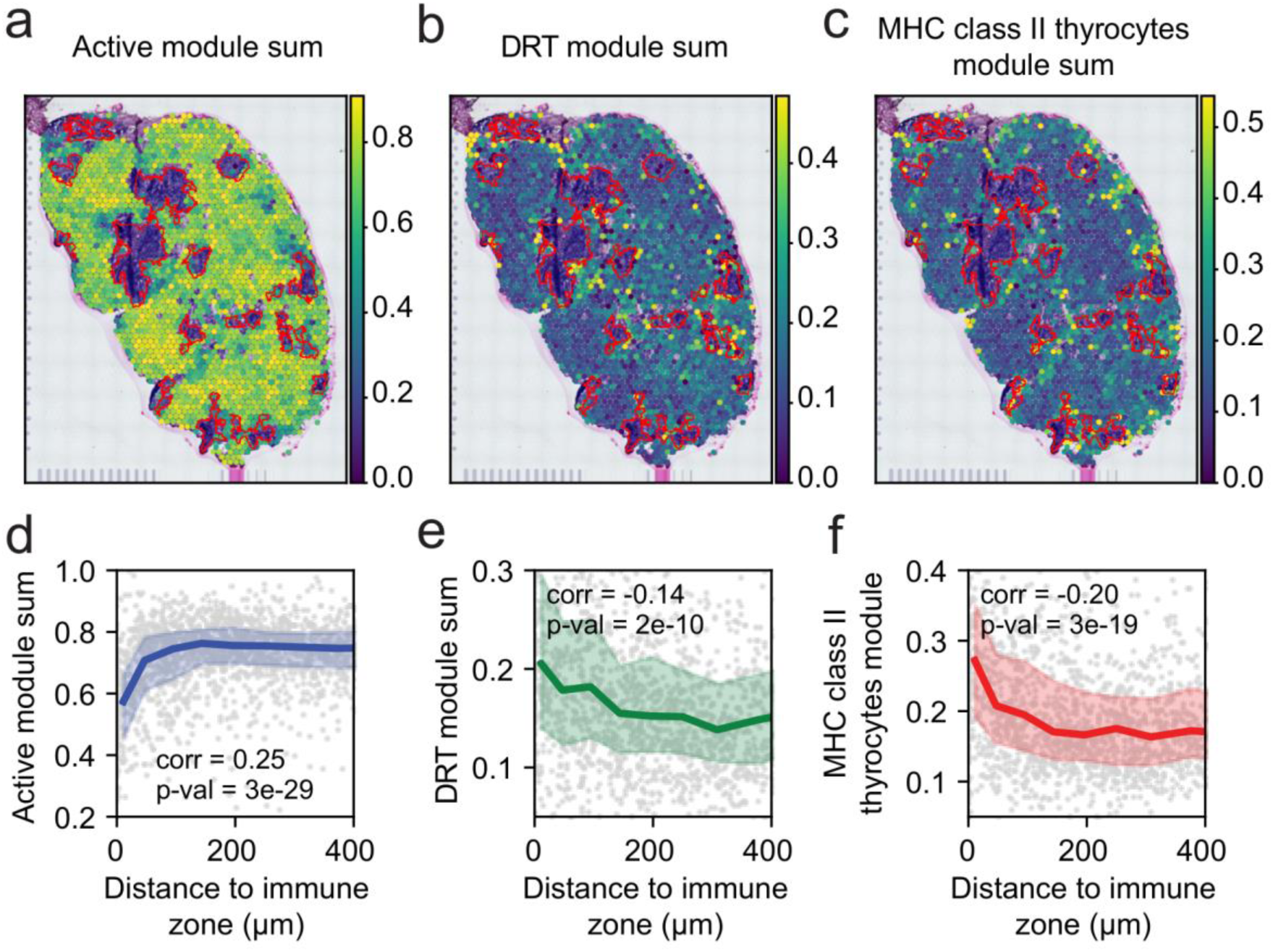
Damage-response thyrocytes are located near immune zones and are distinct from MHC class II⁺ thyrocytes. **a–c**, Spatial maps of module scores for active thyrocytes (**a**), damage-response thyrocytes (DRTs, **b**), and MHC class II thyrocytes (**c**). Lymphocyte aggregates are outlined in red. See also Supplementary Fig. 5a bottom panel. **d–f**, Association between module score and distance to immune zones for active, DRT, and MHC class II programs. Dots represent Visium spots in a representative patient (P9, see additional patient, P8, in Supplementary Fig. 5); curves show median expression in equal-sized bins, with shaded areas indicating the 25^th^-75^th^ percentiles.

Previous studies of inflamed human thyroid tissue have described antigen-presenting thyrocytes which express MHC class II molecules^14,15,27^. In our spatial data, these MHC class II thyrocytes were also significantly enriched near immune zones in inflamed samples (*r*=-0.2, p-value=3×10^−19^, Fig. 4c,f, Supplementary Fig. 5d,g). To examine the relationship between damage-response thyrocytes and MHC class II thyrocytes, we analyzed a published single-cell RNA-sequencing dataset from seven human thyroid samples^15^. We found that the DRT and the active populations were present across this dataset (Supplementary Fig. 6a-c), segregated in distinct clusters (Supplementary Fig. 6d). Moreover, the two modules were strongly anti-correlated across individual cells (Supplementary Fig. 6e-g). MHC class II-expressing thyrocytes covered a distinct transcriptional cluster (Supplementary Fig. 6d-f,h). Comparison of marker gene expression across clusters demonstrated that DRTs were enriched for their marker genes *ARC* and *IL1RL1*, separating them from the active clusters which strongly expressed thyroid hormone synthesis genes such as *TG* and *TPO*, and from the MHC class II thyrocytes, which preferentially expressed antigen-presentation genes including *CD74* (Supplementary Fig. 6e, Table S4). Importantly, the MHC class II thyrocytes were not significantly correlated with DRTs (Spearman *r*=0.007, p-value=0.3, Supplementary Fig. 6f). Together, these analyses indicate that although damage-response thyrocytes and MHC class II-expressing thyrocytes are enriched near immune zones in the thyroid gland, they represent transcriptionally distinct epithelial states rather than overlapping populations.

### Damage-response programs define a major axis of inter-patient variability

To assess whether the balance between active and DRT programs extends beyond spatially resolved measurements, we analyzed bulk RNA sequencing data of human thyroid glands from GTEX^28^ (Methods). Archetype analysis^9,29,30^ revealed three major transcriptional states corresponding to inflamed, active, and damage-response programs (Fig. 5a,b).

**Figure 5.**
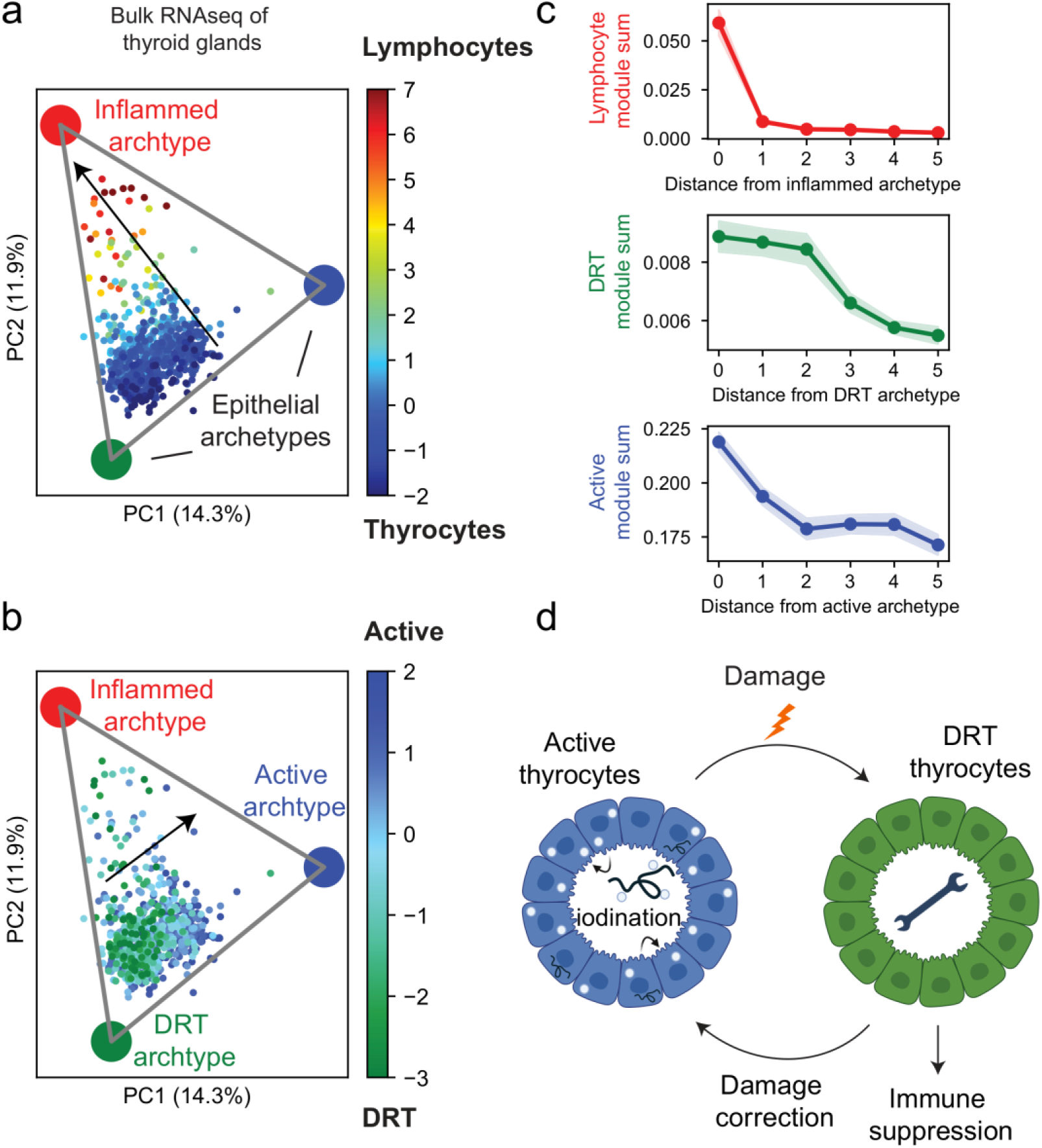
Damage-response programs define a major axis of variability between patients. **a,b,** Principal component analysis (PCA) of bulk RNA-seq profiles from human thyroid glands (n = 684; GTEx^28^), with archetype analysis identifying inflamed, active, and damage-response (DRT) transcriptional states^30^. In **a**, samples are colored by the relative z-score difference between lymphocyte and epithelial gene module expression, showing variability in samples’ lymphocyte content. In **b**, samples are colored by the relative z-score difference between active and DRT gene module expression. **c**, Module behavior as a function of distance from the corresponding archetype (shown in b). Cells were grouped into equal-sized bins by archetype distance; points indicate mean summed gene expression per bin, with shaded regions denoting standard error of the mean. **d**, Conceptual model proposing a division of labor between active thyrocytes and damage-response thyrocytes in the context of damage and damage correction, with hypothesized immune-modulatory consequences.

Distances from these archetypes were associated with module-specific gene expression patterns, with damage-response gene expression defining a dominant axis of variability across samples (Fig. 5c). We further examined the correlations between the DRT and active module gene set expression with patients’ age across the 684 GTEX samples. We found that aging is associated with a significant increase in DRT gene expression and a decrease in active thyrocyte genes (Supplementary Fig. 7a-d). This age-related decline in thyrocyte activity has been demonstrated in a recent scRNAseq study of the human thyroid gland^15^. Notably, we found that the DRT module gene expression significantly increases with age in individual thyrocytes in this dataset as well (Supplementary Fig. 7e-f). These findings indicate that damage-response thyrocytes represent not only a spatially organized epithelial state within tissues but also a major contributor to inter-patient and age-related transcriptional variability.

Based on these findings, we propose a model in which DRTs support adaptation to the metabolic, oxidative, and immune-mediated stresses imposed by sustained hormone production. It has been suggested that excessive oxidative stress may lead to thyroid pathologies^25^. By enabling autonomous stress management at the cellular level, DRTs may limit the escalation to overt immune activation and thereby contribute to the maintenance of functional homeostasis within thyroid follicles (Fig. 5d).

## Discussion

An outstanding question in tissue biology is how cellular heterogeneity is organized within organs composed of repeating anatomical units. In the thyroid gland, one might expect heterogeneity to be organized primarily around the biochemical steps of thyroid hormone synthesis itself, given the complexity and energetic cost of iodide uptake, organification, coupling, and hormone release^31,32^. Alternatively, heterogeneity could reflect stochastic variation or be driven by immune and stress-related cues arising from the tissue microenvironment. Using a spatial analysis that explicitly distinguishes intra-follicular from inter-follicular variability, our study reveals that thyrocyte heterogeneity is not primarily organized by differences in hormone synthesis programs, but instead by stress-response and immune-associated transcriptional states.

Across spatial scales, we identify a damage-response thyrocyte state that emerges as a dominant axis of variability within follicles, between follicles, and across patients. Notably, this state is present even in histologically normal thyroid tissue, suggesting that it represents a physiological epithelial state rather than a purely pathological response. At the same time, in inflamed tissues, DRTs preferentially localize near immune infiltrates, suggesting that this state is dynamically engaged to cope with immune-mediated stress. Together, these observations support a model in which thyroid follicles achieve functional balance through division of labor between hormonally active thyrocytes and damage-repairing cells. Future work utilizing ex-vivo follicle cultures^33–35^ could explore whether the DRT state is irreversible or instead represents a transient, reversible adaptation, and if so, what are the timescales and cues for switching between DRT and active states.

Several lines of evidence suggest that DRTs are engaged in managing oxidative and genotoxic stress associated with sustained hormone production. Thyroid hormone synthesis requires the generation of reactive oxygen species, creating a chronically stressful biochemical environment for follicular epithelial cells. Consistent with this, DRTs are enriched for oxidative stress, p53-family, and DNA damage-response pathways, and exhibit elevated levels of γH2AX. This pattern is consistent with a homeostatic stress-management program that allows follicles to sustain hormone production while limiting cumulative damage. The spatial clustering of DRTs within follicles further suggests that stress responses are locally coordinated, rather than uniformly distributed across all thyrocytes.

Beyond managing intrinsic stress, these observations raise the question of how damage-response thyrocytes interact with the immune system and whether they actively modulate immune responses in the thyroid microenvironment. Our findings point to a potential immunoregulatory role for DRTs through engagement of the IL-33 – IL1RL1 signaling axis. DRTs express *IL1RL1*, the canonical receptor for IL-33, a cytokine with well-established roles in epithelial repair and immune regulation in multiple tissues, including the thyroid (Supplementary Fig. 2). IL1RL1 exists in both a membrane-bound signaling form (ST2L) and a soluble decoy form (sST2) that sequesters IL-33 and blocks downstream signaling^36^. In epithelial contexts, sST2 produced by primary epithelial cells has been shown to act as a negative regulator on inflammation in diseases such as asthma and ulcerative colitis^37–39^. This raises the possibility that DRTs locally suppress IL-33-mediated immune activation by producing soluble ST2, thereby limiting immune recruitment while engaging autonomous epithelial repair programs. Disruption of this regulatory circuit could contribute to persistent inflammation and may help explain the association between *IL1RL1* genetic variants and autoimmune thyroid disease^40^.

An unexpected feature of the damage-response thyrocyte program is the prominent expression of *ARC*, a domesticated retrotransposon-derived gene best characterized in neurons^21^. In the nervous system, ARC assembles into virus-like capsids that mediate intercellular RNA transfer and are released from neurons, often within extracellular vesicles, enabling molecular communication between cells. Although ARC has not previously been implicated in thyroid biology, its expression in DRTs raises the possibility that analogous intercellular communication mechanisms could operate in epithelial tissues. Such mechanisms may contribute to the clustered spatial organization of DRTs we observed within and between follicles, suggesting that stress-adapted states may propagate locally rather than arising independently in each cell. Notably, other domesticated retrotransposon-derived, capsid-forming proteins expressed in the central nervous system, such as PNMA2, have been proposed to be immunogenic when ectopically expressed^41^. This observation points to a broader class of capsid-based signaling mechanisms that may acquire immunogenicity when dysregulated.

More broadly, our study provides a conceptual and analytical framework for dissecting heterogeneity in tissues composed of repeating anatomical units. For example, in a multi-tasking organ such as the liver, hepatocytes perform hundreds of functions and operate in concentric layers along liver lobules, repeating hexagonal units^42^. One could envision division of labor between hepatocytes that reside within the same lobule layer, or sub-specialization of distinct lobules for different tasks. Similar approaches could be applied to the intestinal villi, the kidney nephrons and other structured organs, where cells must balance multiple, sometimes competing functions under spatially patterned metabolic and immune constraints.

In summary, our work demonstrates that stress-response and active epithelial states dominate the organization of thyrocyte heterogeneity across spatial scales in the human thyroid gland. The identification of a damage-response thyrocyte state that is both physiological and dynamically engaged in immune contexts provides insights into how the thyroid gland could maintain functional integrity in the face of metabolic and immunological stress and illustrates how multiscale spatial analysis can uncover organizing principles in complex tissues.

## Data availability

The spatial transcriptomics datasets generated in this study will be publicly available upon publication via Zenodo at https://doi.org/10.5281/zenodo.18494459.

## Code availability

Scripts will be available in GitHub upon publication.

## Competing interests

The authors declare that they have no competing interests.

## Acknowledgements

We thank Dan Ryan and Prof. Jason Shepperd for discussions.

The R.M. laboratory is supported by the Howard Hughes Medical Institute, Food Allergy Science Initiative, Blavatnik Family Foundation, and Colton Center for Autoimmunity. SI is supported by the Moross Integrated Cancer Center, the Helen and Martin Kimmel Award for Innovative Investigation, the Yad Abraham Research Center for Cancer Diagnostics and Therapy, the Israel Science Foundation MAVRI program grant no. 1482/25, the European Union (ERC, GI-DYNAMICS, 101198168), the Swiss Society Institute for Cancer Prevention Research, and a Weizmann-Schneider joint research grant and a research grant from the Ministry of Innovation, Science and Technology, Israel. Optical imaging data was acquired at the de Picciotto Cancer Cell Observatory In memory of Wolfgang and Ruth Lesser of the Moross Integrated Cancer Center in the department of Life Science Core Facilities, Weizmann institute of science. YKK is supported by the JSMF Postdoctoral Fellowship in Understanding Dynamic and Multi-scale Systems (Award #https://doi.org/10.37717/2020-1428).

---

**Supplementary Figure 1.**
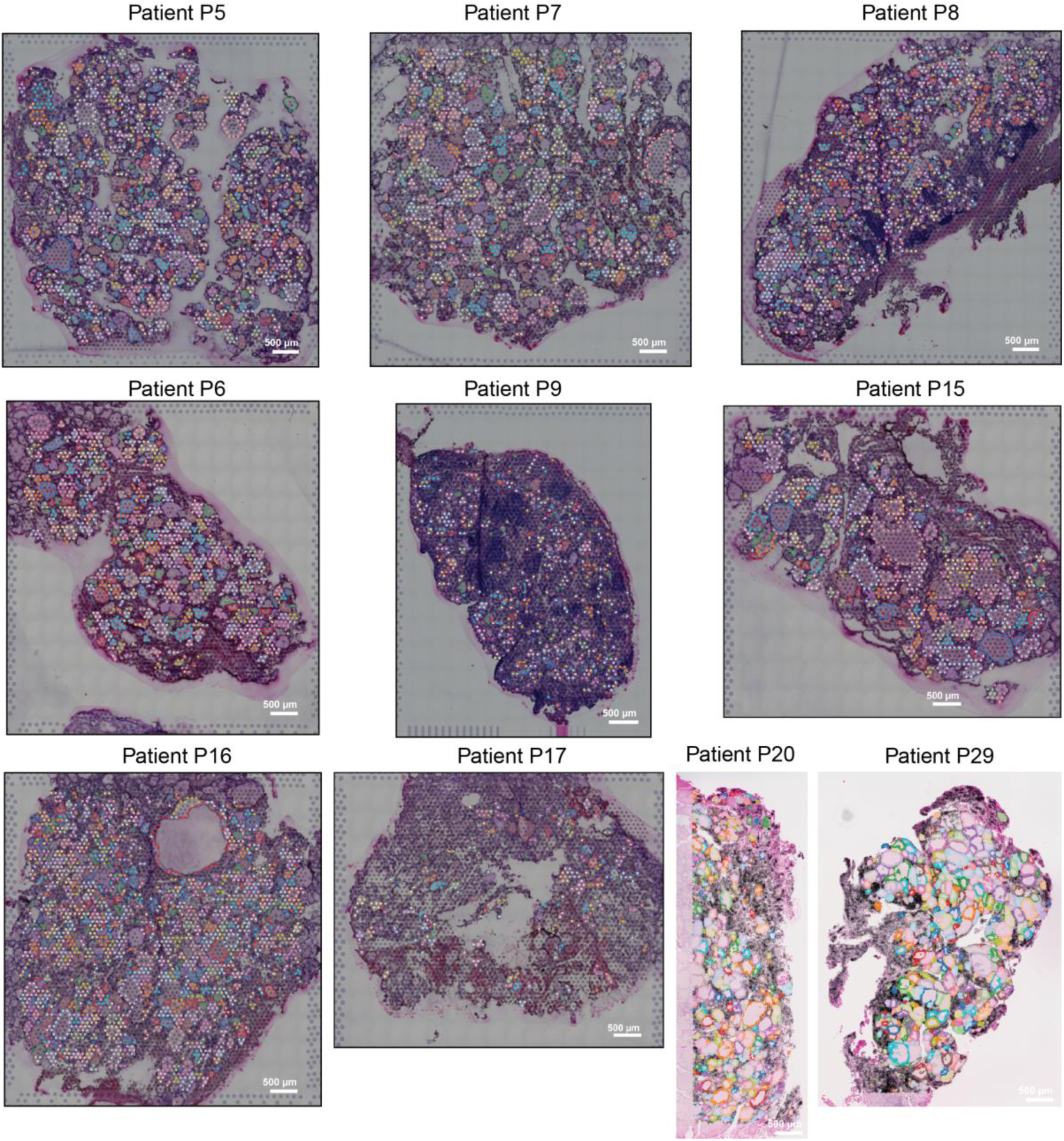
Follicle segmentation across human thyroid samples. H&E images showing Visium spots (P5-P17) and VisiumHD reconstructed cells (P20, P29) overlaid with follicle segmentation. Spots/cells are randomly colored according to the follicles they are assigned to, with unannotated spots/cells in gray.

**Supplementary Figure 2.**
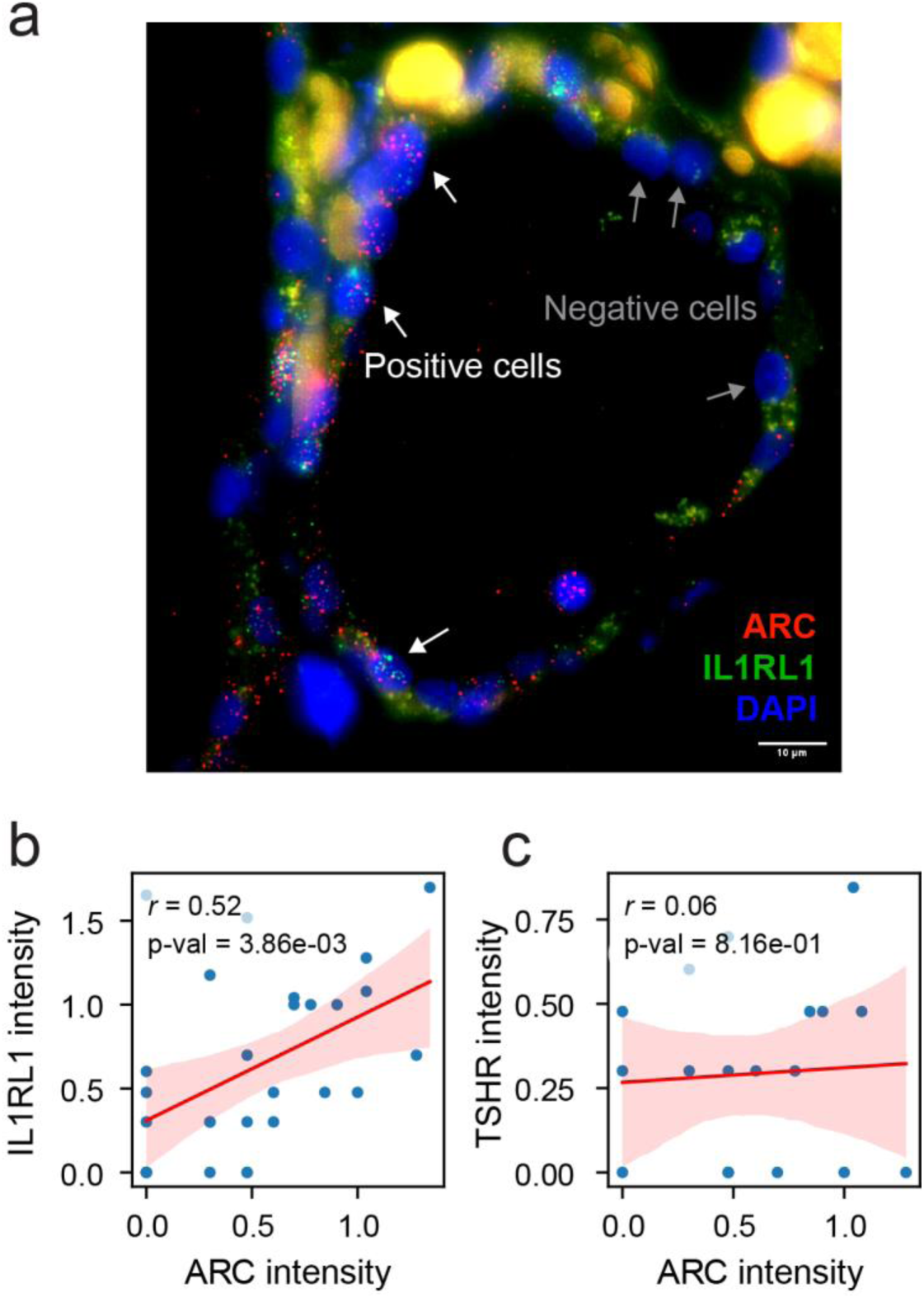
ARC and IL1RL1 are co-expressed. **a,** Representative HCR FISH image of a thyroid follicle showing expression of *ARC* (red) and *IL1RL1* (green) in thyrocytes, with nuclei stained by DAPI (blue). White and gray arrows indicate positive or negative cells for *ARC* and *IL1RL1* expression, respectively. Scalebar 10µm. **b,** Quantification of HCR FISH signal intensity in single cells demonstrating a positive correlation between *ARC* and *IL1RL1* expression across individual thyrocytes in the representative image. Spearman correlation denoted by r. **c,** Lack of correlation between *ARC* and *TSHR* expression across the same cells (negative control), indicating specificity of *ARC–IL1RL1* co-expression.

**Supplementary Figure 3.**
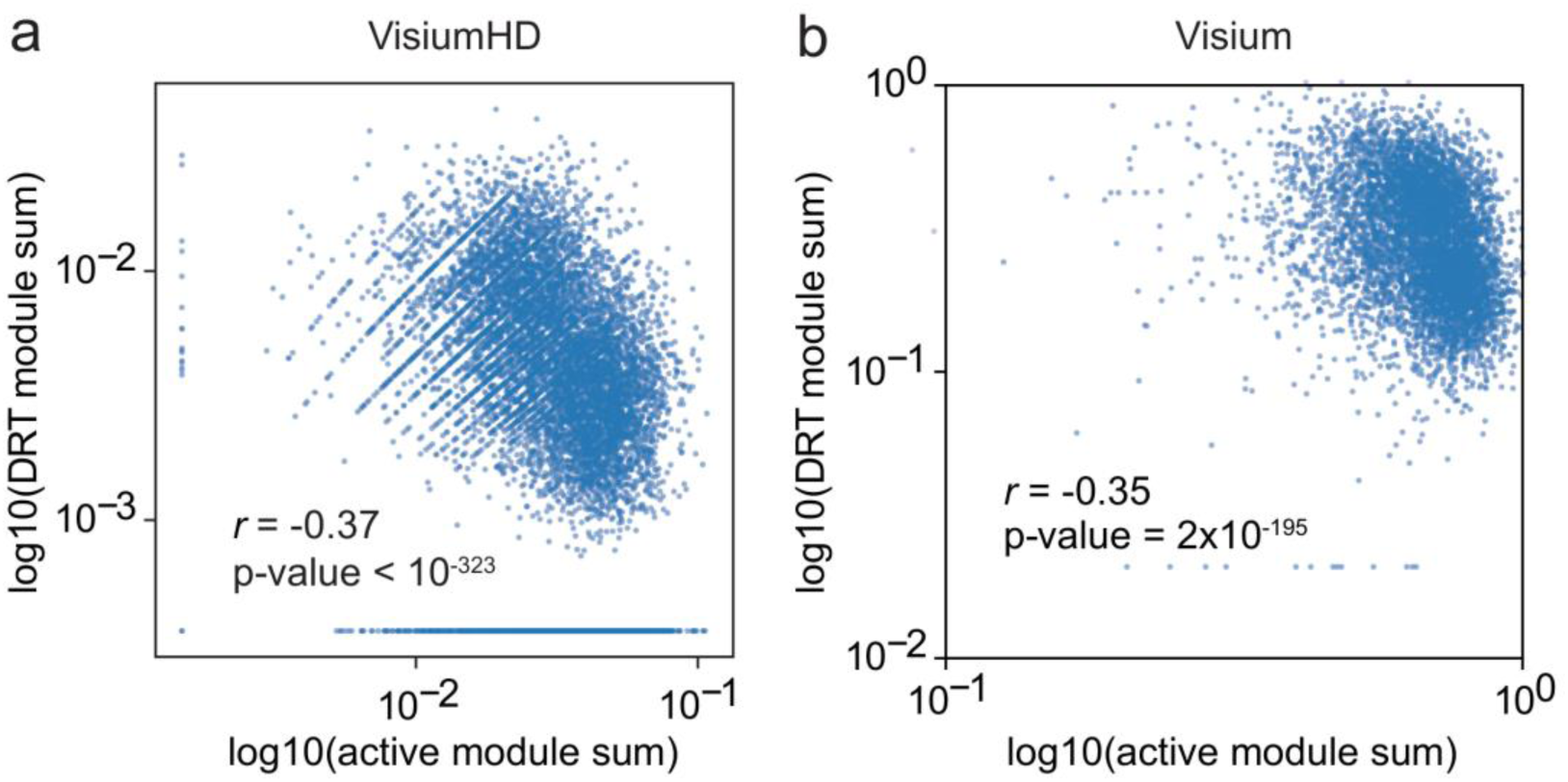
Active and damage-response modules are anti-correlated. Scatter plots showing the relationship between active thyrocyte module scores and DRT module scores at the cellular level in VisiumHD data (**a**) and spot level in Visium data (**b**). Spearman correlation coefficients and p-values are shown. For Visium spots, module sums were normalized to total thyrocyte module sum.

**Supplementary Figure 4.**
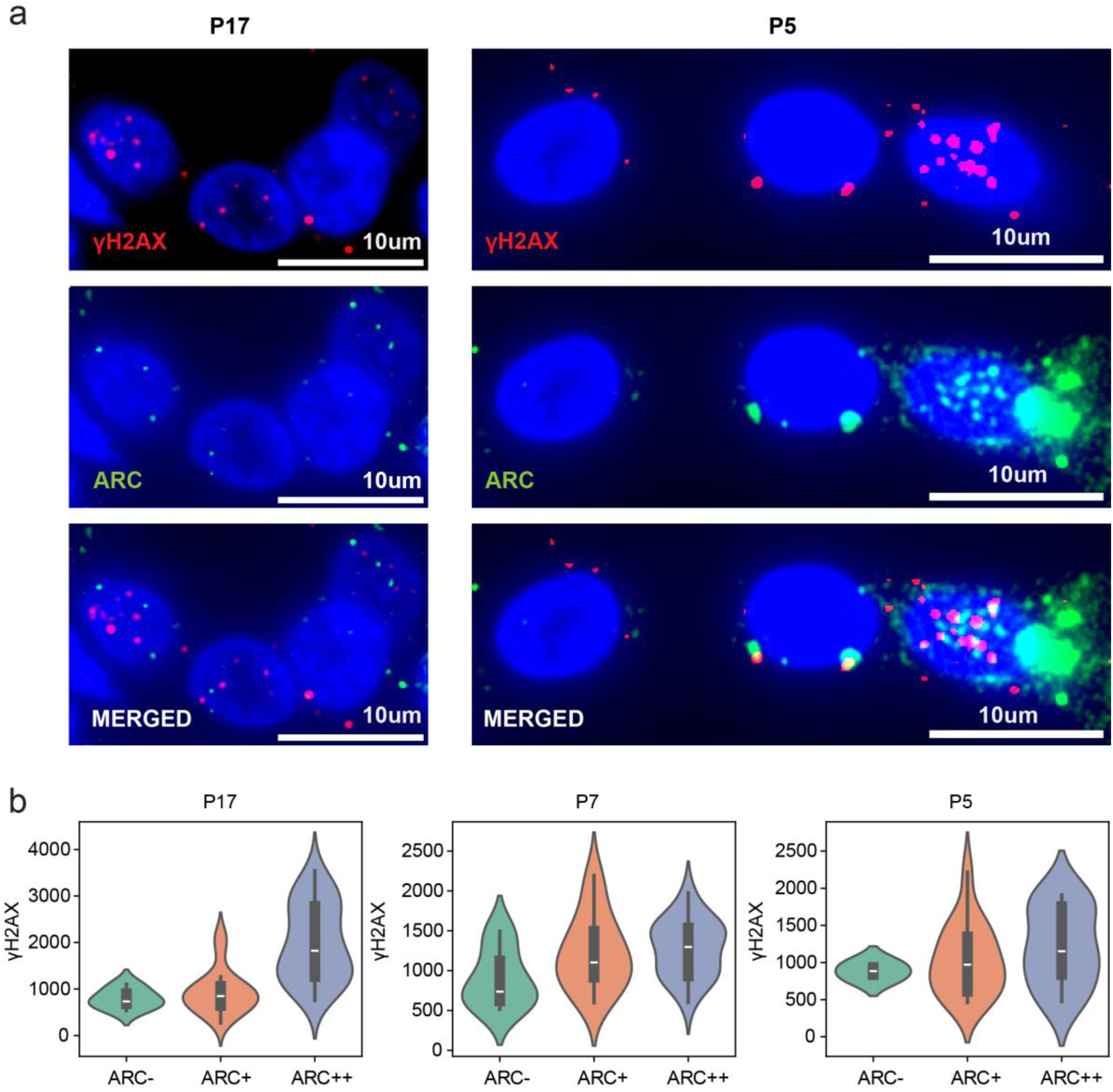
Association between *ARC* expression and γH2AX signal across patients. **a,** Representative zoomed-in images of combined HCR FISH and immunofluorescence (IF) staining showing expression of γH2AX protein (red) and *ARC* mRNA (green) in thyrocytes, with nuclei stained by DAPI (blue). **b,** Violin plots showing γH2AX fluorescence intensity in ARC-negative, ARC-positive, and ARC-high thyrocytes for individual patients (Methods). Increased γH2AX signal is consistently observed with higher *ARC* expression, supporting association of the damage-response thyrocyte state with DNA damage response. Box plots white line indicates the median, boxes span the IQR (25,75 percentiles), whiskers extend up to 1.5 IQR.

**Supplementary Figure 5.**
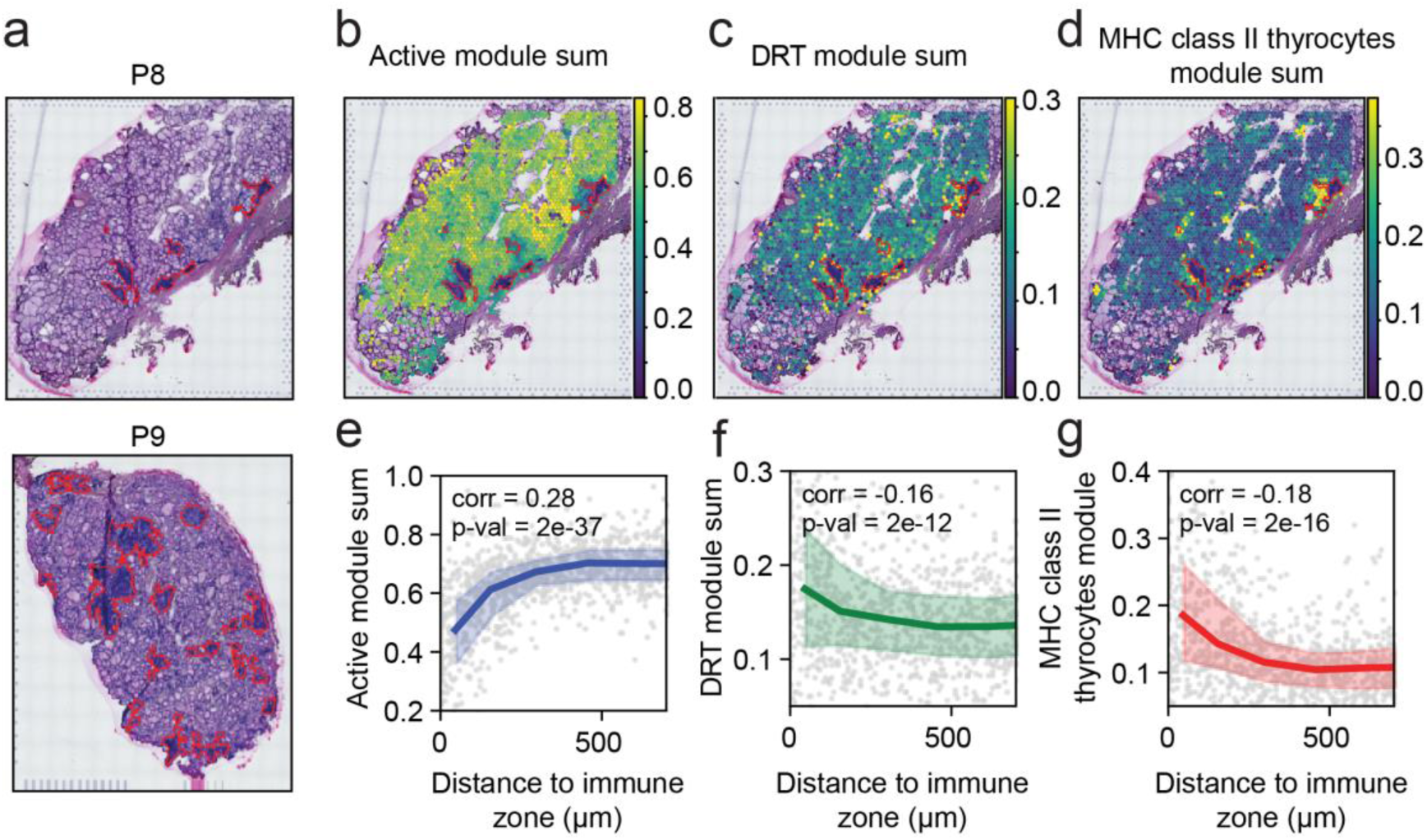
Damage-response thyrocytes are located near immune zones and are distinct from MHC class II⁺ thyrocytes. **a** H&E staining with lymphocyte aggregates outlined in red for patient P8 (top panel) and patient P9 (Bottom panel, see Fig. 4). **a–c**, Spatial maps of module scores for active thyrocytes (**a**), damage-response thyrocytes (DRTs, **b**), and MHC class II thyrocytes (**c**) for P8. Lymphocyte aggregates are outlined in red. **d–f**, Association between module score and distance to immune zones for active, DRT, and MHC class II programs in P8. Dots represent Visium spots; curves show median expression in equal-sized bins, with shaded areas indicating the 25^th^-75^th^ percentiles.

**Supplementary Figure 6.**
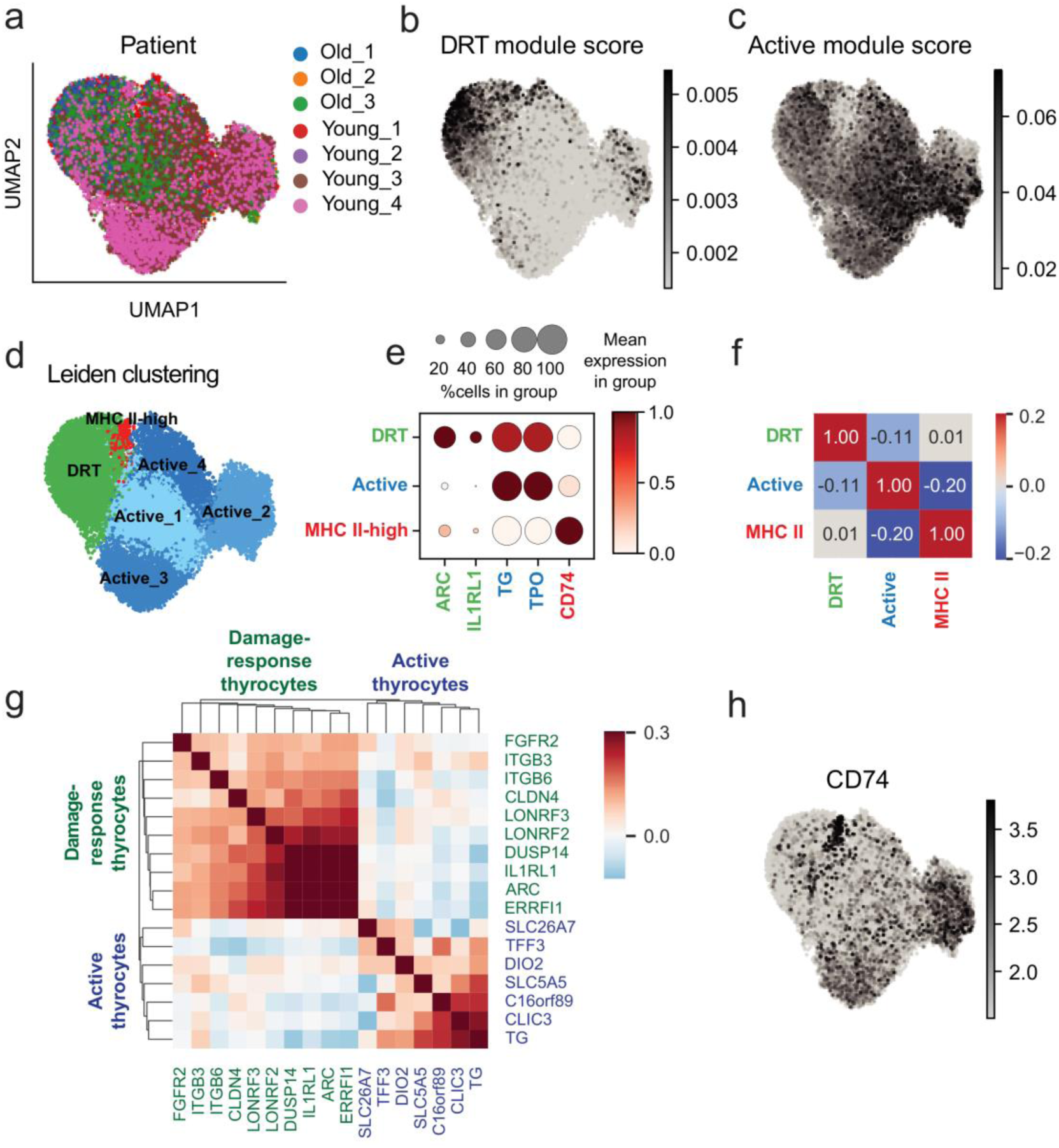
Damage-response and active thyrocyte transcriptional programs are present in single-cell data and are distinct from MHC class II thyrocytes. **a,** UMAP embedding of thyrocytes from a published single-cell RNA-sequencing dataset of human thyroid samples^15^, colored by patient. **b,c,** Projection of DRT module scores **(b)** and active module scores **(c)** onto the UMAP, showing broad representation of both programs across cells. Module scores are the summed gene expression of the module genes, normalized to sum of thyroid-specific genes. **d,** Leiden clustering of thyrocytes identifies DRT cluster, multiple active clusters, and a transcriptionally distinct MHC class II cluster. **e,** Dot plot showing expression of selected marker genes across clusters; dot size indicates the fraction of cells expressing each gene and color indicates scaled mean expression level. **f,** Spearman correlation coefficients between DRT, active and MHC class II modules in single cells. **g,** Heatmap showing pairwise Spearman correlations between selected marker genes across single cells. **h,** UMAP projection of log-transformed *CD74* expression.

**Supplementary Figure 7.**
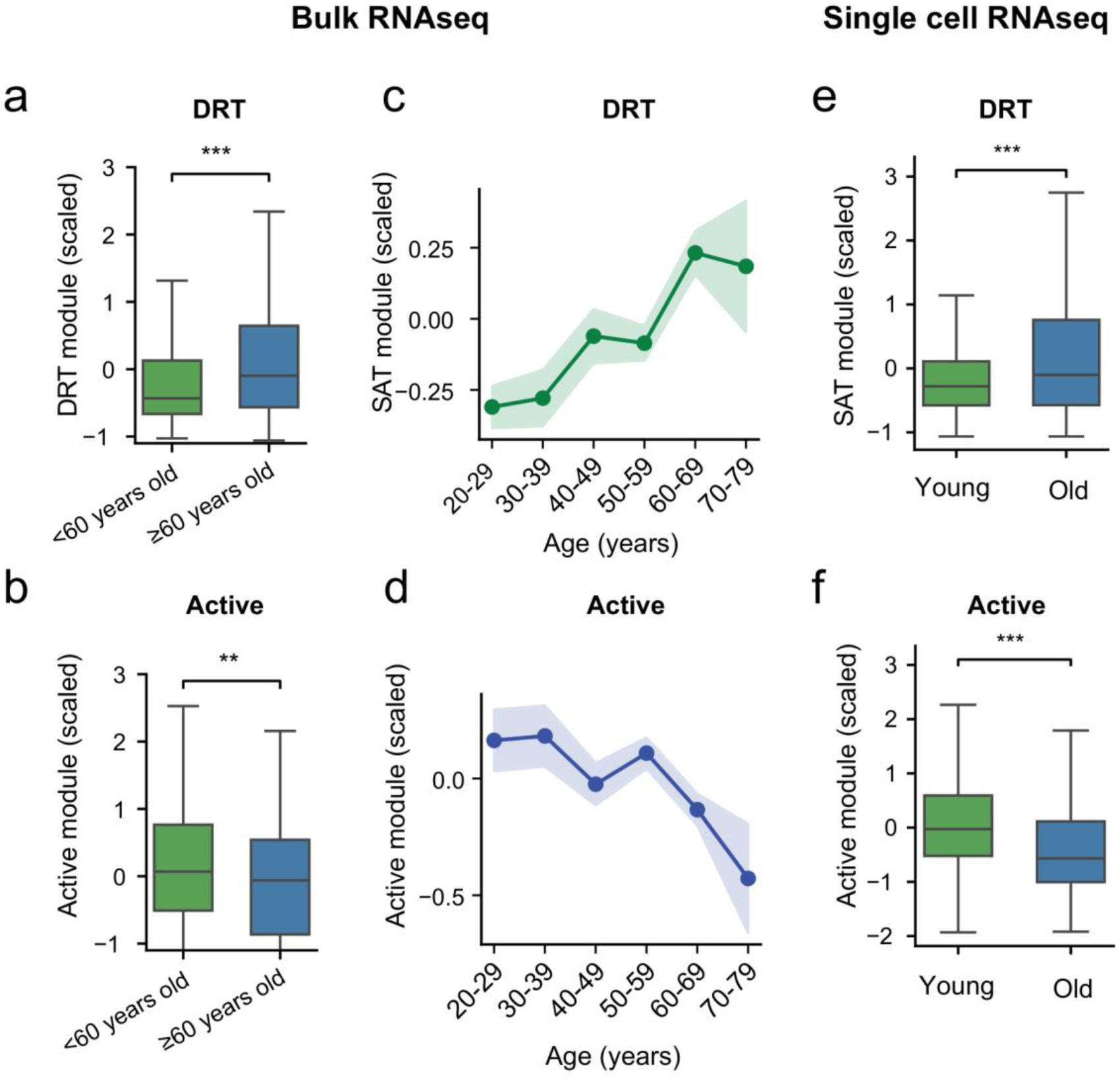
Age-associated shift between damage-response and active thyrocyte states. **a,b,** Comparison of DRT **(a)** and active **(b)** Module scores between younger (<60 years) and older (≥60 years) individuals in bulk RNA-seq data of human thyroid glands (GTEx^28^). Box plots indicate median and interquartile range (25,75 percentiles), with statistical significance indicated (p-values: 4e-06 (DRT), 8e-3 (active)). Module scores were scaled by z-score normalization. **c,d,** Scaled DRT module scores **(c)** and active thyrocyte module scores **(d)** shown across age decades in bulk RNA-seq data. Shaded areas are standard errors of the mean. **e,f,** Scaled DRT **(e)** and active **(f)** module scores derived from single-cell RNA-sequencing data^15^, comparing thyrocytes from younger (n=4, 20-30 years old) and older (n=3, 50-60 years old) donors. Box plots indicate median and IQR (25,75 percentiles), whiskers extend up to 1.5 IQR. Statistical significance is indicated (p-values: 1e-55 (DRT), 2e-238 (active)).

**Supplementary Table 1 | Metadata for human thyroid samples analyzed in this study**

Clinical and technical metadata for all human thyroid samples included in the study, including sample identifiers, patient demographics, histological assessment, immune infiltration status, and spatial transcriptomics platform used.

**Supplementary Table 2 | Intra-follicular gene expression variability**

Results of the intra-follicular variability analysis for thyrocyte-specific genes passing expression and inclusion thresholds (see Methods). Variability was quantified across thyrocytes within individual follicles and compared to a multinomial randomization-based null model. Mean follicle std denotes the mean standard deviation of UMI counts in follicles in which the gene was detected. Mean z-score indicates the mean standardized deviation relative to the null distribution. p-values were obtained by combining follicle-level p-values, and q-values represent false discovery rate-adjusted p-values. # follicles indicates the number of follicles contributing to the gene-level statistics.

**Supplementary Table 3 | Inter-follicular gene expression variability**

Results of the inter-follicular variability analysis for thyrocyte-specific genes passing expression and inclusion thresholds (see Methods). Mean patient std is the mean standard deviation of expression across follicles averaged over patients. Mean z-score is the mean deviation from random expectation across patients. p-values were obtained by combining patient-level p-values, and q-values represent false discovery rate-adjusted p-values.

**Supplementary Table 4 | Gene expression in DRT, active and MHC class II modules based on single-cell gene expression data**

Signature matrix of DRT, active and MHC class II thyrocyte modules, from single cell data of human thyroid glands^15^. Clusters and selected genes are shown in Supplementary Fig. 5d,e. Expression is given as number of UMIs normalized to total UMI counts in each cell. Genes with expression lower than 10^−6^ normalized UMI count were excluded.

## Methods

### Patient cohort and tissue processing

Patients undergoing surgical procedures were recruited from the Department of Otolaryngology at Sheba Medical Center under an approved institutional review board protocol (Helsinki approval no. SMC-9569-22). Written informed consent was obtained from all participants prior to inclusion in the study. The cohort included individuals undergoing total thyroidectomy, hemithyroidectomy, or total laryngectomy (n = 10; 5 male and 5 female). Clinical and demographic information was obtained from electronic medical records (Table S1).

Immediately following surgical excision, the resected thyroid specimen was examined by a senior surgeon, and a small tissue fragment was dissected from a region distant from the pathological area. Depending on tissue volume, specimens were subdivided into up to three portions for downstream applications: (i) fresh-frozen embedding in optimal cutting temperature (OCT) compound (Scigen, 4586) followed by storage at −80 °C; (ii) formalin fixation and paraffin embedding (FFPE; fixation in 4% formaldehyde (J.T. Baker, JT2106) for 24 h, followed by 1% formaldehyde for an additional 24 h, and subsequent processing at the Weizmann Institute of Science Histology Unit); and (iii) formalin fixation (4% formaldehyde for 3 h, followed by overnight incubation in 4% formaldehyde with 30% sucrose), cryo-embedding in OCT, and storage at −80 °C.

### Visium

Visium (10X Genomics) was carried out according to manufacturer instructions. Briefly, fresh frozen OCT embedded tissues were sectioned into 10μm thick and placed on the Visium spatial gene expression slide (10X genomics). Tissues were fixed with methanol and stained with H&E. Brightfield images were taken on a Leica Widefield-DMI8 microscope with 20X magnification. Tissues were permeabilized for 16-20 minutes, based on a Visium spatial tissue optimization user guide (10x Genomics, CG000238). Libraries were prepared according to the Visium Spatial Gene Expression user guide (CG000239). Libraries were quantified using TapeStation analysis (Agilent) and NEBNext library prep kit for Illumina (NEB), and sequenced on the Novaseq 6000 using SP 100 cycles kit. Further processing was done using 10X Space Ranger software 1.3.0.

### VisiumHD

10X VisiumHD was performed on thyroid gland samples from 2 patients (Table S1). Sections (5 µm thick) from FFPE tissue blocks were mounted onto VisiumHD–compatible slides and processed for hematoxylin and eosin (H&E) staining according to the 10X VisiumHD FFPE Tissue Preparation Handbook (CG000684). Brightfield whole-slide images were acquired using a Leica DMi8 widefield inverted microscope equipped with a Leica DFC7000T color camera and an HC PL APO 20×/0.80 dry objective (506530; Leica Microsystems CMS GmbH). Following imaging, tissue sections were processed for spatial transcriptomics using the VisiumHD Spatial Gene Expression Reagent Kits (CG000685) and CytAssist system (v2.1.0.14) at 37 °C for 30 min, in accordance with the manufacturer’s instructions. Generated libraries were quantified using TapeStation analysis (Agilent) and NEBNext library prep kit for Illumina (NEB) and loaded onto individual lanes of NovaSeq X Plus (Illumina) using a 1.5B 100 cycles flow cell. Further processing was done using 10X Space Ranger software 3.0.0.

### HCR FISH

*In-situ* HCR was performed according to the Molecular Instruments HCR protocol for FFPE tissues. In brief, FFPE sections were deparaffinized and rehydrated through a graded ethanol series (Xylene, Xylene, 100% Ethanol, 100% Ethanol, 95% Ethanol, 70% Ethanol, 50% Ethanol, PBS). Sections were permeabilized for 10 min with proteinase K (10 mg ml−1 in PBS, Ambion AM2546) followed by two washes with PBS. Tissues were incubated in hybridization buffer (HCR hybridization buffer with 100 μg ml−1 salmon sperm DNA, Sigma-Aldrich D7656) for 10 min in a 37 °C incubator. Tissues were incubated with hybridization buffer mixed with 0.4 pmol probes for 16 h in 37 °C. After the hybridization, tissues were washed with decreasing concentrations of HCR wash buffer in 5× SSCT (5× SSC, 0.1% Tween-20) for 15 min in 37 °C. The washes were performed successively with buffer concentrations of 100%, 75%, 50%, 25% and, finally, 0%. The samples were washed with 5× SSCT for 5 min at room temperature, then incubated with AB (HCR amplification buffer with 100 μg ml−1 salmon sperm DNA) for 30 min at room temperature. HCR-hairpins (H1 and H2 hairpins of B1-546, B3-594, B5-647) were heated for 90 s at 95 °C and cooled in dark for 30 min. The samples were incubated with AB with 6 pmol of each hairpin for 45 min in the dark, and then rinsed in 5× SSCT for 5, 15 and 15 min. The samples were incubated with 5× SSCT with DAPI (50 ng ml−1, Sigma-Aldrich, D9542) for 5 min. Additional wash with 5× SSCT was performed, and slides were mounted with Immu-Mount (Epredia, 990402). Images were taken with 100x magnification on the Nikon-Ti-E inverted fluorescence microscope using the NIS element software AR v.5.11.01.

### HCR FISH combined with HCR immunofluorescence

*In situ* HCR combined with immunofluorescence was performed according to the Molecular Instruments HCR protocol for FFPE tissues. In brief, FFPE sections were deparaffinized and rehydrated through a graded ethanol series (Xylene, Xylene, 100% Ethanol, 100% Ethanol, 95% Ethanol, 70% Ethanol, 50% Ethanol, PBS). Sections were permeabilized for 10 min with TritonX (0.5% in PBS) followed by two washes with PBS. Tissues were incubated in hybridization buffer (HCR hybridization buffer with 100 μg ml−1 salmon sperm DNA, Sigma-Aldrich D7656) for 10 min in a 37 °C incubator. Tissues were incubated with hybridization buffer mixed with 0.4 pmol probes overnight in 37 °C. After the hybridization, tissues were washed with decreasing concentrations of HCR wash buffer in 5× SSCT (5× SSC, 0.1% Tween-20) for 15 min in 37 °C. The washes were performed successively with buffer concentrations of 100%, 75%, 50%, 25% and, finally, 0%. The samples were washed with 5× SSCT for 5 min at room temperature, then blocked with HCR antibody buffer with 5% normal horse serum for 1 hour at room temperature. Samples were incubated with rabbit anti Phospho-Histone-H2A.X antibody (1:100, Cell signaling #2577) overnight at 4 °C. Samples were washed 3 times with PBS for 5 minutes and incubated in antibody buffer with 5% normal horse serum with anti-rabbit-B5 secondary antibody (1:500) for 1 hour at room temperature. Samples were washed 3 times with PBS for 5 minutes and then fixed with 4% PFA for 10 minutes. Samples were washed 3 times with PBS and incubated with AB (HCR amplification buffer with 100 μg ml−1 salmon sperm DNA) for 30 min at room temperature. HCR-hairpins (H1 and H2 hairpins of B1-546, B3-594, B5-647) were heated for 90 s at 95 °C and cooled in dark for 30 min. The samples were incubated with AB with 6 pmol of each hairpin for 45 min in the dark, and then rinsed in 5× SSCT for 5, 15 and 15 min. The samples were incubated with 5× SSCT with DAPI (50 ng ml−1, Sigma-Aldrich, D9542) for 5 min. Additional wash with 5× SSCT was performed, and slides were mounted with ProLong Gold Antifade Mountant (Invitrogen P36930). Images were taken with 60x magnification, on the Nikon-Ti-E inverted fluorescence microscope using the NIS element software AR v.5.11.00.

### γH2AX-*ARC* co-expression analysis

Nuclei were segmented using Stardist algorithm in Fiji^43,44^. γH2AX mean intensity was measured for each nucleus. ARC low, intermediate and high cells were detected by a blinded observer. Analysis included 7 images from 3 different patients (Supplementary Fig. 4). p-value was computed by Kruskal-Wallis *H*-test on the pooled cells from all patients.

### Histological segmentation and spatial assignment

Follicles were segmented from H&E-stained tissue images using the Cellpose algorithm^45^ implemented in QuPath v0.5.0^46^, with parameters optimized to identify follicles rather than cells (model=”cyto2”, diameter=100). Segmented follicle boundaries were expanded by 1%, and all segmentations were manually reviewed to remove misannotated regions. For VisiumHD, cell nuclei were segmented from H&E-stained tissue images using the StarDist algorithm implemented in QuPath v0.5.0. Segmented nuclei were expanded by up to 10 µm or until contacting a neighboring cell to approximate full cell boundaries. VisiumHD bins were assigned to individual cells when the bin centroid fell within the corresponding cell polygon. Gene expression counts from all bins assigned to a cell were summed to generate the final cell-level gene expression matrix.

To generate follicle-level gene expression matrices, cells were assigned to follicles if their centroid fell inside the corresponding follicle polygon. For Visium data, spots were assigned to follicles based on centroid inclusion, and spots in the follicular lumen, defined as spots with all six neighboring spots located within the same follicle, were removed (Fig S1). Immune zones (Fig. 4), defined as lymphocyte-rich regions, were segmented using a supervised pixel classifier trained in QuPath v0.5.0. A random trees pixel classifier at moderate resolution was trained using all available multiscale image features and scales. Objects were then created from immune class pixels using a minimum object and hole size of 30000µm^2^.

### Defining thyrocyte-specific genes

Thyrocyte-specific genes were defined based on signature matrix from published human thyroid single-cell RNAseq data (Hong et al.^15^), containing epithelial cells, endothelial cells, fibroblasts, myeloid cells, proliferating cells, SMCs/Pericytes, B cells and T/NK cells. Thyrocyte-specific genes were defined as genes above a minimal expression threshold (5×10^−6^, with total cell expression normalized to 1), which are at least 2-fold higher in thyrocytes than in any other cell type.

### Cell type annotation of VisiumHD data

To define cell types in the cell-level VisiumHD data, we applied Cell2location algorithm^47^ using the reference annotations from Hong et al.^15^, and assuming one cell per spatial location. Cells were assigned to the cell type with the highest posterior probability (epithelial cells, endothelial cells, fibroblasts, myeloid cells, smooth muscle cells/pericytes, proliferating cells, T/NK cells, and B cells).

### Intra-follicular variability analysis

Intra-follicular variability was quantified using a multinomial randomization framework, to compare to a null model which preserves total UMIs per cell and per gene within each follicle. Analyses were restricted to thyrocytes located within segmented follicles, as defined by spatial annotation, and to thyrocyte-specific genes (Methods) expressed above a minimum mean expression threshold (5×10^−6^ sum normalized expression). Cells with fewer than 20 total UMIs (out of thyrocyte-specific genes only) and follicles containing fewer than three thyrocytes were excluded from analysis.

For each gene and follicle, variability was quantified as the standard deviation (SD) of UMI counts across thyrocytes within the follicle. To generate a null distribution, gene-specific UMI counts within each follicle were randomly redistributed across cells using multinomial sampling. This procedure was repeated 100 times to estimate the expected mean and SD of intra-follicular variability under the null model. Genes were retained for analysis only if variability statistics could be computed in more than 10 follicles.

Observed SD values were converted to *z*-scores relative to the null distribution and transformed into one-sided p-values using the survival function of the standard normal distribution. For each gene, p-values were combined across follicles using Pearson’s method, and genes represented in fewer than three follicles were excluded. Multiple hypothesis testing correction was performed using the Benjamini–Hochberg procedure to control the false discovery rate.

### Inter-follicular variability analysis

Inter-follicular variability was quantified to assess gene expression heterogeneity across follicles within each patient using a multinomial randomization framework. Analyses were restricted to thyrocyte-specific genes (Methods).

For each patient, gene expression was aggregated at the follicle level by summing UMI counts across spots (for Visium data) or cells (for VisiumHD) within each follicle. Inter-follicular variability for each gene was quantified as the standard deviation of summed UMI counts across follicles within a patient. Spots and cells with ≤20 total UMIs (summed across the thyrocyte marker genes) were excluded. Follicles containing fewer than 3 Visium spots were excluded. Genes with ≤30 total thyrocyte-specific gene UMIs summed across follicles for a given patient were excluded.

To generate a null distribution, total UMI counts for each gene were randomly redistributed across follicles using multinomial sampling, with probabilities proportional to the total UMI content of each follicle, thereby preserving per-gene total counts and follicle-level sequencing depth. This randomization procedure was repeated 100 times to estimate the expected mean and standard deviation of inter-follicular variability under the null model.

Observed inter-follicular standard deviations were converted to z scores relative to the null distribution and transformed into one-sided p-values using the survival function of the standard normal distribution. Gene-level p-values were computed independently for each patient and then combined across patients using Fisher’s method, requiring p-values from at least two patients. Multiple hypothesis testing correction was performed using the Benjamini–Hochberg procedure to control the false discovery rate.

### Pathway enrichment analysis on DRT and active module genes

Pathway enrichment analysis was performed using the Enrichr^20^ API implemented in GSEApy^48^ (Python) against the MSigDB Hallmark 2020 and KEGG 2021 Human gene sets, using thyroid-specific genes as the background set. Gene modules used for pathway enrichment analysis were defined by hierarchical clustering of the gene-gene correlation matrix constructed from genes exhibiting significant inter-follicular variability (Benjamini-Hochberg adjusted q-value < 0.05; Table S2). The heatmap shown in Fig. 2d represents a subset of this correlation matrix (q-value<10^−5.4^). The resulting dendrogram was cut at a maximum distance threshold of 0.985, yielding two gene modules comprising 51 genes for the DRT module and 213 genes for the active thyrocyte module.

### Spatial auto-correlation analysis

Follicles were embedded in a spatial coordinate system by computing their centroids, and Euclidean distances were used to identify each follicle’s nearest spatial neighbor. DRT and active cellular module scores were computed and z-scored, as in *spatial module plots*, and their difference was used to classify cells as DRT-positive or DRT-negative (z-score(active) – z-score(DRT)<0). Spatial concordance was defined as the fraction of neighboring follicle pairs sharing the same module identity. To assess whether concordance exceeded random expectations, module identities were permuted across follicles while preserving the neighbor graph, and concordance was recalculated across 1000 permutations to generate a null distribution.

An analogous analysis was performed at the single-cell level. Cells were assigned to follicles, and each cell’s nearest neighbor was determined only among cells within the same follicle, ensuring that spatial relationships reflected local organization rather than global tissue structure. Follicles were included in this analysis only if they contained at least ten module-positive cells to ensure adequate statistical power. Cells were classified using an analogous module-score thresholding strategy, and concordance between neighboring cells was quantified as described for follicles. A permutation-based null model was generated by shuffling module identities within each follicle 1000 times while keeping intra-follicular neighbor relationships fixed, allowing comparison of observed cell-level concordance to randomized expectations.

### Bulk GTEX samples analysis

Bulk RNAseq thyroid samples gene expression and metadata were acquired from the GTEX database^28^ (n=684, 225 females, 459 males). Lowly expressed genes with mean expression less than 1 TPM across samples were removed. PCA was performed on the log-normalized data, and archetypes were found using the PCHA algorithm with delta=0.7.

In Fig. 5a,b, each gene module was summed, z-scored, and cells are colored by the z-score difference of the respective modules (lymphocyte-specific and epithelial-specific gene modules in 5a, DRT and active modules in 5b). Lymphocyte and epithelial specific genes were defined based on annotated single cell data from Hong et al^15^. Cell type specific genes were defined as genes with cell type expression at least 2-fold higher than in any of the other cell types. DRT and active gene modules were defined based on the intra-follicular variability analysis (see heatmap in Fig. 2d).

## Notes

### Competing Interest Statement

The authors have declared no competing interest.

